# Effect of Nivolumab therapy on Metastatic Lung Cancer in Human Microbiome

**DOI:** 10.1101/2023.01.03.522679

**Authors:** Rohan Kubba, Robert J. Evans, Sonha Nguyen, Ramadas Pai, James Borneman

**Affiliations:** UCLA B.S. (Biology) 2023 Minor in Global Health; FACG; FRCP, FACC, UCR Medical School; UCR Department of Plant Pathology and Microbiology

## Abstract

Nivolumab, a type of immunotherapy, has enhanced the 5-year survival of patients with renal cell cancer, melanoma, and lung cancer which is now mechanistically understood. However, relatively sparse information assesses its relationship with shaping the gut microbiome. We aimed to assess the degree to which nivolumab treatment contributes to alterations in the species composition of the colon in lung cancer patients undergoing nivolumab treatment. Our pilot study utilized stool samples of five lung cancer patients at Inland Hematology Oncology (IHO) before administration of nivolumab and three months after initiation of treatment. 16S and ITS rRNA sequence analysis were used to assess alterations in species abundance and richness of the colon. After sequencing, statistical analysis, specifically a paired t-test, was performed to assess if any significant differences in any microbial species were observed before and after immunotherapy treatment. Although different proportions of microorganisms existed at baseline prior to treatment for each patient, a significant reduction in the *Megasphaera elsdenii* population was observed (p=.0488; n=4), when comparing before and after treatment. Our findings differ from that of Huang et. al (2022), who has recently posited that a positive association exists between *Megasphaera elsdenii* and the survival of patients with pancreatic ductal adenocarcinomas. Our conclusions suggest that different cancers may elicit differential effects on *Megasphaera elsdenii* in the gut microbiome.

**Importance:** Nivolumab is a relatively new form of cancer therapy called immunotherapy, which enables the body to use its own immune system to effectively fight cancer. Due to advances in microbial sequencing in 16S rRNA, this project explores differences in the abundance of microbial communities before and three months of treatment in patients who have advanced on chemotherapy. Our findings suggest that reduction of *Megasphaera elsdenii*, a metabolically active bacterium, is associated with positive outcomes, which differs from findings from other literature. Our project advocates for a more robust profiling of the microbiome during lung cancer treatment, and immunotherapy, in particular, to establish a more substantive profile of the changing gut in the midst of treatment.

## Background

Cytotoxic T cells, (CD 8 + cells) recognize cancer cells initially using major histocompatibility complex I (MHC I); however before eradication, cancer cells employ PD-L1 (programmed cell death ligand 1), a checkpoint inhibitor, on its surface to bind to PD-1 (programmed cell death protein 1), which subsequently downregulates cytotoxicity. Cancer cells effectively become undetectable to the immune system, hijacking the PD-1/PDL-1 pathway. Nivolumab (Opdiv™) capitalizes on this pathway as its mechanism of action consists of binding its antibody to the PD-1 receptor of a cytotoxic T-cell, effectively blocking the cancer cells from interfering with T lymphocyte cytotoxicity, thereby inducing an onslaught of cytokines and proteolytic factors that promote cancer cell death.

Recent literature has suggested that immunotherapy treatment in both gastric cancer (Lau et al. 2020) and melanoma (Davar et al. 2021) has led to alterations in the gastrointestinal (GI) microbiome composition. Additionally, Carbone et al. (2019) explained that symbiotic associations of microorganisms accentuated the efficacy of the immune system in lung cancer response. Our pilot study aimed to study the effects of nivolumab on host stool microbial communities and to look for associations with alterations in microbiota and tumor responsiveness in lung cancer patients.

## Methodology

Patients with advanced stage lung cancer refractory to chemotherapy presenting to Inland Hematology Oncology consented to provide a stool sample for microbial analysis prior to and three months after initiation of therapy with nivolumab. The gut bacteria were identified using an illumina-based rRNA ITS and 16S rRNA (ribosomal ribonucleic acid internal transcribed spacer) sequence analysis; the specific product is denoted as OMNIgene®•GUT from DNA GenoTek. 16S rRNA sequencing allowed us to assess relative prokaryotic abundance and richness, and quantify differences in proportions at the three month period. Baseline computed tomography (CT) scans were obtained and repeated at a three month interval, established in the protocol, to determine tumor status. Overall, patients 1, 2, 4 and 5 had stable disease or improvement with nivolumab according to their scans. The stool analysis primarily focused on multiple groups of species (19) and genera (19).

All samples were sequenced by Dr. Borneman of the Department of Plant Pathology and Microbiology at the University of California Riverside. Patients with adenocarcinoma showed improved responses. Unfortunately, patient 3 showed progression of disease and had squamous cell cancer histology.

As evidenced by Figures 1 and 2, patient 3 endured a complete restructuring of the gut microbial community, which correlated with unfavorable outcome. Moreover, patient 3 had drastic variances in the colon microbiota ultimately harboring over sixty percent of *Bacteroides theta iota omicron* species in an overwhelming relative abundance.

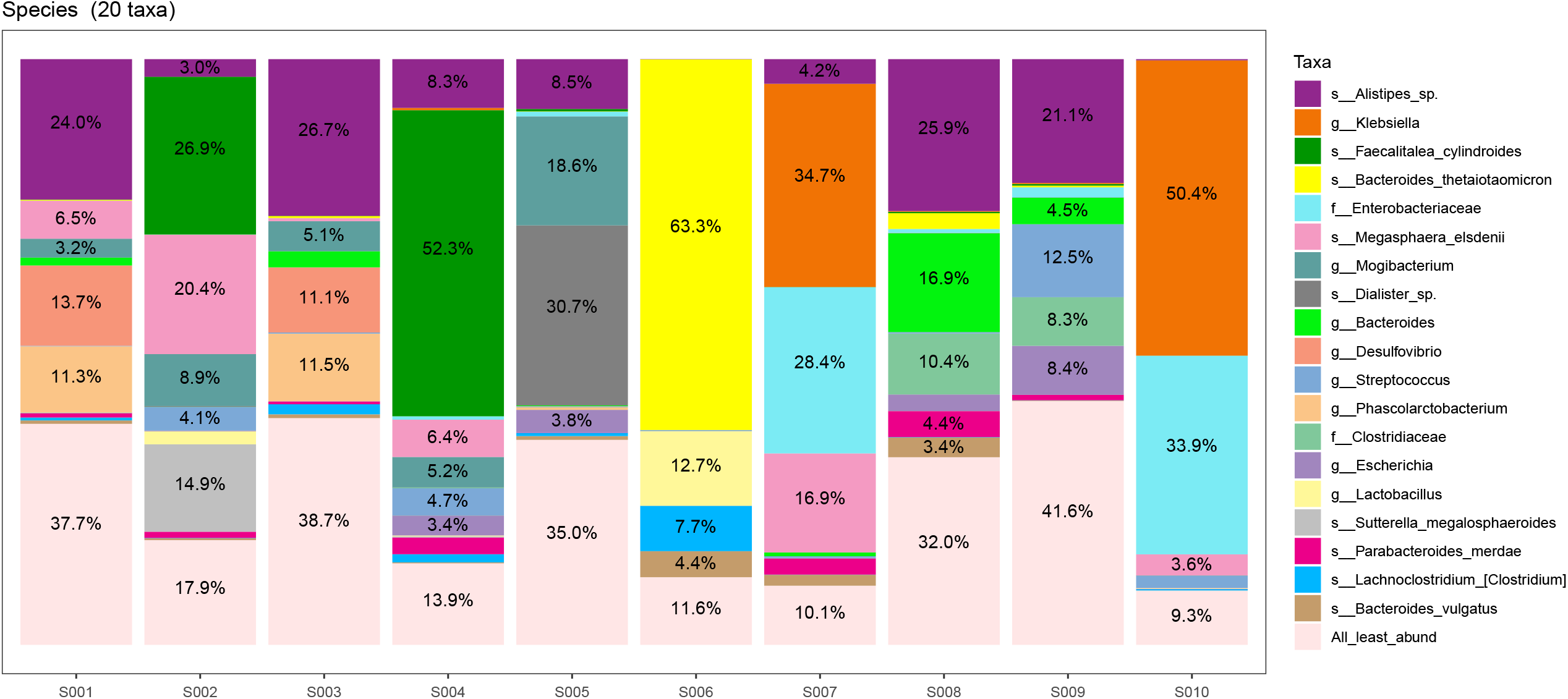

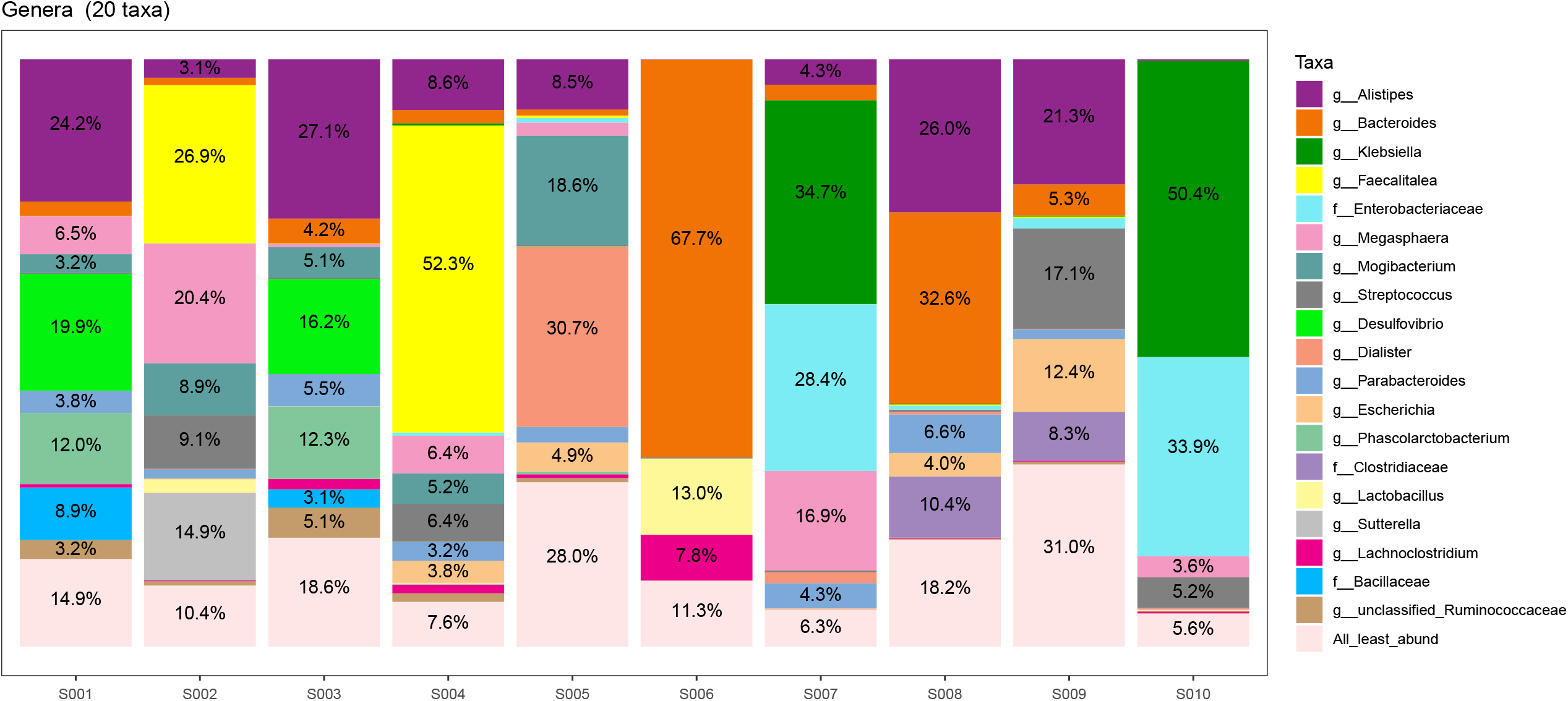

When assessing the effect of nivolumab *Megasphaera elsdenii* on the subjects, data from patient 5 was omitted. Patient 5 did not have any proportion of *Megasphaera e*. colonization prior to treatment and also did not gain abundance. Additionally, patients 1, 2, and 4 demonstrated a significant difference in their *Megaspheare e*. values before and after nivolumab treatment (p=.042, n=3). Although patient 3 began with a sub-5% level of *Megasphaera e*., when accounted for in the t-test, the results would still be statistically significant (p=.0488, n=4)..

### Data Availability Statement

Bacterial ITS rRNA sequencing data was deposited in the National Center for Biotechnology Information (NCBI) Sequence Read Archive under BioProject accession number: PRJNA900790. There are ten total samples in this entry, corresponding to the before-treatment and after-treatment for each of the 5 patients.

## Discussion

Shetty et al. (2013) contend that the bacterium *Megasphaera elsdenii* possesses functional genes that potentially synthesize essential amino acids and vitamins as well as enhance the breakdown of carbohydrates in the human host. They cite that *Megasphaera elsdenii* is a chief contributor to the synthesis of short chain fatty acids (SCFAs), such as butyrate, which is a preferred energy source in the colon and also functions to bolster the immunity function of the cell membrane by maintaining pH levels and compartmentalization. SCFAs are also known inhibitors of histone deacetylases in colon cells, which consequently upregulate transcriptional activity in the gastrointestinal tract. Huang et al. (2022) indicated that an increase in *Megasphaera elsdenii* is positively correlated with better health outcomes, using murine models and quantifying the level of tumor growth inhibition. They found that the presence of *Megasphaera elsdenii* with an anti-PD-1 checkpoint inhibitor led to decreased cancer growth than the checkpoint inhibitor alone. Our data suggests that differential cancer types and treatment may have conflicting impacts on the gut microbial community.

Cancer is not a monolith and different cancers have the ability to express variable effects on the gut microbiome. A possible hypothesis for the discrepancy between our findings and that of Huang et al. (2022) may be due to differences in cancer variety, eliciting differing results physiologically, or sampling bias as the patients in Huang et al (2022) were primarily of Chinese ethnicity while our study was primarily Hispanic. Although characterized by seemingly beneficial interactions with their host, it has been associated with positive outcomes in pancreatic/colorectal cancers. However, our findings suggest that attenuation of *Megasphaera e*. is associated with better outcomes in lung cancer patients.

A common side effect of nivolumab treatment is inflammation of the gut likely disrupting the distribution of microbial species. Recruitment of a variety of macrophages, natural killer cells, and other leukocytes may all be mechanisms of action for modulating species richness; however further research exploring this association should be more extensive and be more globally encompassing.

While our data reveals a plethora regarding alterations in gastrointestinal microbial population information with the usage of five patients, it is necessary to conduct this research on a much larger scale, potentially minimizing confounding variables In order to substantiate our conclusions regarding the interaction of immunotherapy and lung cancer, it is necessary to survey on normalized populations as well. Our project primarily focuses on a group with a median age of 62. It is important to note that this report only scratches the surface of the potential data that the gut microbiome holds in terms of different cancers and their interaction with therapy that can potentially affect outcomes. Therefore, we advocate for further research that continues to track the gut microbiome in cancer patients to ultimately further advancements in the field of microbiology and cancer understanding.

## Acknowledgments

All funding, specifically the stool kits, was done by Inland Hematology Oncology with no external conflicts of interest. To reiterate, there is no pharmaceutical or industry funding involved at all with this project. Dr. Borneman ran all of the samples at no cost in his laboratory at the University of California Riverside. Additionally, Alexander Pang of the Department of Statistics at the University of California Los Angeles performed a paired t-test to determine statistical significance of *Megasphaera elsdenii* percentage before and after treatment.

## Note

Opdivo™ is made by Bristol-Myers Squibb Company, a pharmaceutical company headquartered in the USA on 430 E. 29^th^ Street, 14^th^ Floor, New York, NY 10016.

OMNIgene®·GUT is made by DNA Genotek(r) at 500 Palladium Dr #3000, Kanata, ON K2V 1C2, Canada

## Legend for Figures 1 and 2

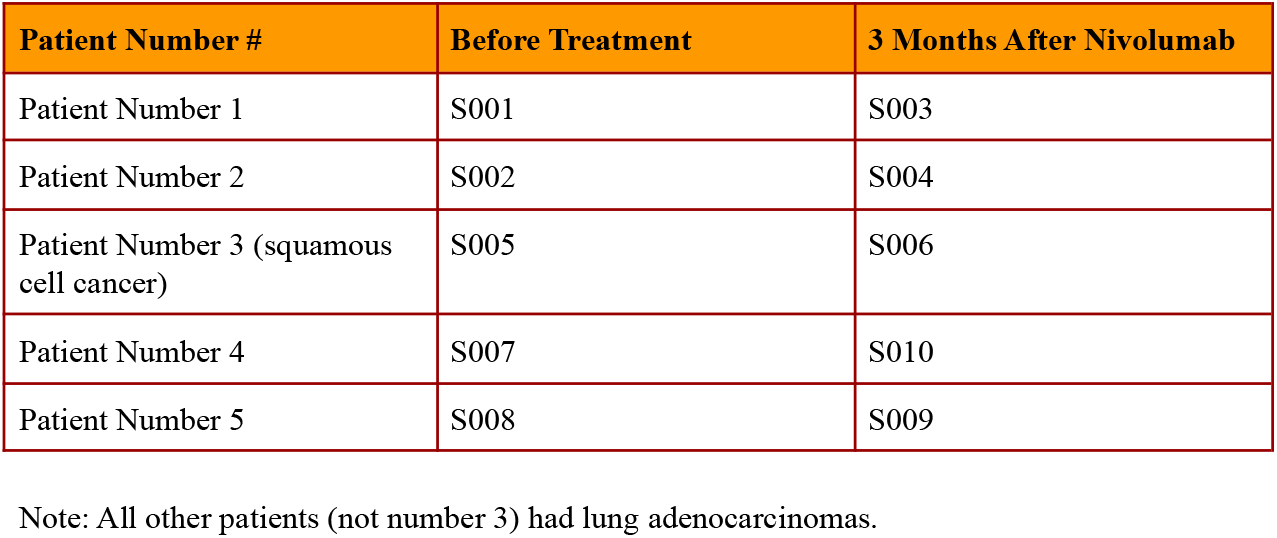

| Patient Number # | Before Treatment | 3 Months After Nivolumab |
| --- | --- | --- |
| Patient Number 1 | S001 | S003 |
| Patient Number 2 | S002 | S004 |
| Patient Number 3 (squamous cell cancer) | S005 | S006 |
| Patient Number 4 | S007 | S010 |
| Patient Number 5 | S008 | S009 |

Note: All other patients (not number 3) had lung adenocarcinomas.

